# IsoAnalytics: A Single-cell Proteomics Web Server

**DOI:** 10.1101/2023.01.03.522673

**Authors:** Suzette N. Palmer, Andrew Y. Koh, Xiaowei Zhan

## Abstract

**Motivation:** Single-cell proteomics provide unprecedented resolution to examine biological processes. Customized data analysis and facile data visualization are crucial for scientific discovery. Further, userfriendly data analysis and visualization software that is easily accessible for the general scientific community is essential.

**Results:** We have created a web server, <u>IsoAnalytics</u>, that gives users without computational or bioinformatics background the ability to directly analyze and interactively visualize data obtained from the Isoplexis single cell technology platform. We envision this open-sourced web server will increase research productivity and serve as a free, competitive alternative for single-cell proteomics research.

**Availability:** IsoAnalytics is free and available at: https://cdc.biohpc.swmed.edu/isoplexis/ and is implemented in Python, with all major browsers supported. Code for IsoAnalytics is free and available at: https://github.com/zhanxw/Isoplexis_Data_Analysis.

**Supplementary Information:** Supplementary data are available at *Bioinformatics* online.

## 1 Introduction

Single-cell proteomics is a high throughput technology that enables quantitative protein profiling of individual cells. The Isoplexis platform utilizes flow-cell and multiplex ELISA technology to allow for the detection of up to 32 cytokines from individual immune cells. This technology serves as an alternative to flow cytometry and mass cytometry (cytometry by flight of time). The Isoplexis technology has provided key insights for a wide variety of biological processes, including cancer, immunology, and oncology (Abbas *et al*., 2021; Baer *et al*., 2022; Kwak *et al*., 2013).

Currently, the only available analysis software for Isoplexis assays is IsoSpeak, which is provided with the machine but offers limited user analyses and options. To offer users greater flexibility and customizability for Isoplexis data analyses, we have created a web server, IsoAnalytics. IsoAnalytics provides expanded data analysis, visualization options and contains four categories of analysis and visualization: Clustering, Dimensionality Reduction, Polyfunctionality, and Statistics. Our web server is completely interactive and does not require any computational or bioinformatics knowledge or coding ability to analyze and visualize data.

## 2 Implementation and Application

IsoAnalytics was created using Dash (Dash, 2022; Plotly, 2015) and can be accessed through https://cdc.biohpc.swmed.edu/isoplexis/. All figures generated by IsoAnalytics are interactive and customizable. For example, zoom and rotation manipulation are available for each visualization. Selection of conditions or variables to be visualized is included. Additionally, all visualizations can be exported as PNG images.

The IsoAnalytics website lands to the “Overview” tab. It gives descriptions for each data analysis provided. Additionally, the cytokines for each Isoplexis single cell secretome assay and the corresponding dominant functional group classifications are provided for easy access. The next tab is “Upload” and allows users to directly load data (CSV or Excel format files) onto the website. The user must then select the correct single-cell secretome assay and then select the appropriate experimental condition or conditions to analyze. The order of this selection will determine the order of these conditions on the graphs. The user also has the option to select which individual cytokine will be analyzed, located on the other analysis tabs. At any time, the user can return to the “Upload” tab and change which treatment conditions and individual cytokines are analyzed.

The data analysis and visualization tabs are Clustering, Dimensionality Reduction, Polyfunctionality, and Statistics. Under “Clustering”, hierarchical clustering across cytokines is performed, and a dendrogram and heatmap are generated for the selected treatment condition(s) **(Fig 1A)**. Underneath this section, for the selected cytokine from the “Upload” tab, hierarchical clustering across cells is performed for samples that have non-zero values for this cytokine, with an accompanying dendrogram and heatmap generated **(Fig 1A)**.

**Fig. 1.**
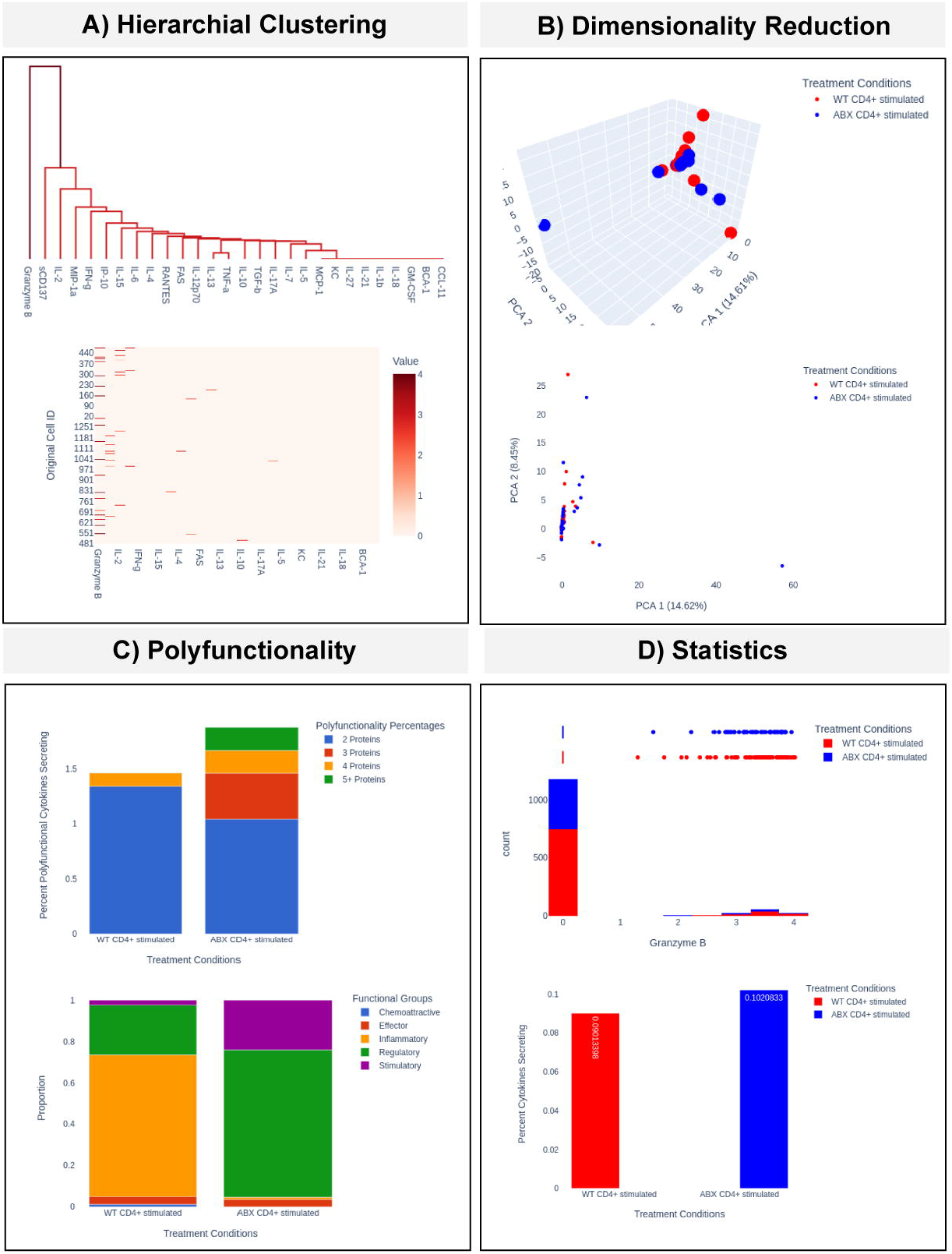
IsoAnalytics webserver data analysis and visualization highlights. **(A)** Hierarchical clustering on Isoplexis data is performed and visualized using a dendrogram (top) and heatmap (bottom). The user has the option to visualize clustering across all provided data or can select to view clustering based on samples containing a specific cytokine. **(B)** Dimensionality reduction analysis PCA can be visualized in 3D (top) or 2D (bottom). **(C)** The stacked bar graph (top) displays the number of polyfunctional cells (cells expressing more than on cytokine) for each treatment condition. The stacked bar graph (bottom) displays the functional groups for each treatment condition. **(D)** The histogram and box plot (top) displays the distribution of the user-selected cytokine and is plotted based on treatment conditions. The user also has the option to visualize the data as a violin or rug plot, instead of the boxplot. The bar plot (bottom) shows non-zero proportions based on the treatment conditions. More detailed descriptions of the analyses and visualizations are available in the IsoAnalytics Supplemental Material.

The “Dimensionality Reduction” tab displays 2D or 3D visualization of Standard Scalar Normalized PCA (Pearson, 1901; Pedregosa *et al*., 2011) **(Fig 1B)** and t-distributed stochastic neighbor embedding (t-SNE) (Van de Maaten and Hinton., 2008; Pedregosa. *et al*., 2011). For the t-SNE algorithm, the user can modify the algorithm by selecting a different perplexity and number of iterations. The “Polyfunctionality” tab displays the number of polyfunctional cells and a stacked bar graph displaying the percent polyfunctional cells for each treatment condition **(Fig 1C)**. The Dominant Functional Groups for each treatment condition are displayed using stacked bar graphs **(Fig 1C)**. The user can select whether to view these data as absolute abundance or as a proportion. Additionally, all data from this tab are available to the user and can be exported as a csv file.

The last tab, “Statistics” displays the Isoplexis data distribution and statistical tests. At the top, non-zero proportions for each treatment condition and cytokine are displayed as an interactive bar graph. The individual cytokine statistics are shown for the selected data or individual treatment conditions. The user can also calculate statistical significance by using the percent cytokines secreting (Non-Zero Proportion) test or the Kolmogorov-Smirnov Test (Virtanen *et al*., 2020) **(Fig 1D)**. The distribution of the data is displayed using a histogram and density plot, where values for each treatment condition are calculated **(Fig 1D)**. Above the histogram, the user can also select whether to view the data as a boxplot, violin plot or a rug plot.

## 3 Results

With the development of novel technology, such as the Isoplexis singlecell functional proteomics platform, new data analysis and visualization tools that are accessible to a wide variety of scientists are essential. We believe IsoAnalytics fills this niche, allowing users without computational or bioinformatics backgrounds to directly and interactively explore data produced by Isoplexis technology. We envision this website will complement the currently available IsoSpeak software, allowing users more options for data analysis and visualization. In future versions, we will include other Isoplexis analyses, such as Codeplex and will continue to add novel assays that are developed for the Isoplexis platform.

## Acknowledgements

The authors would like to thank other members of the Koh lab for crucial feedback that allowed the successful development of the website.

