## Supplementary texts for "IsoAnalytics: A Single-cell Proteomics Web Server"

### Supplemental Material

Sections:


1. Tutorial for IsoAnalytics
    a. Upload Isoplexis Data
    b. Hierarchical Clustering Analysis
    c. Dimensionality Reduction Analysis
    d. Polyfunctionality
    e. Statistics and Distribution
2. FAQs

#### Section 1: Tutorial for IsoAnalytics

1. Login to the following Webserver: https://cdc.biohpc.swmed.edu/isoplexis/. The user will be directed to the "Overview" tab, which contains descriptions of the analysis and visualization pages, details of the Isoplexis single cell secretome assays and contact information.
   b. For issues, please create a new issue through https://github.com/suziepalmer10/Isoplexis_Data_Analysis/issues.

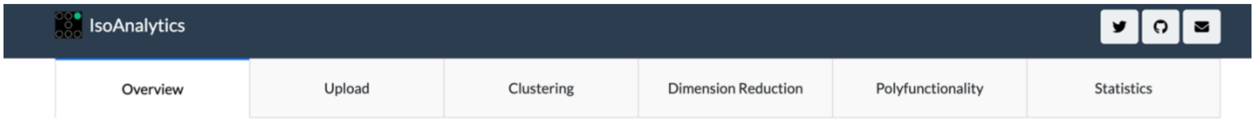


2. To begin the analysis, select the Upload tab. Follow all five steps listed below.
   a. Step 1: Upload Isoplexis single cell data, which can be formatted as a CSV or Excel (.xlsx). Once the file is successfully uploaded, the name of the file and the timestamp of the file will appear below the instructions for step 1.

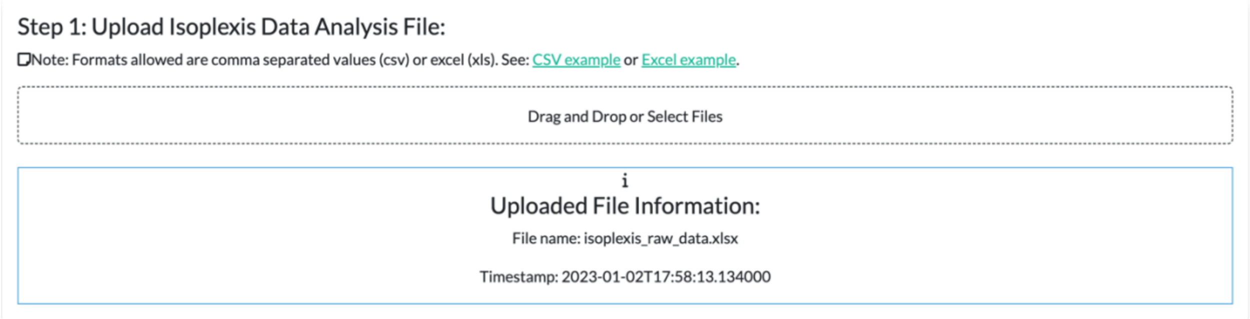


   b. Step 2: Select the assay/panel used for the Isoplexis data. This will continue to be updated as Isoplexis assays are modified and/or developed. Current Isoplexis single cell assays are shown below.

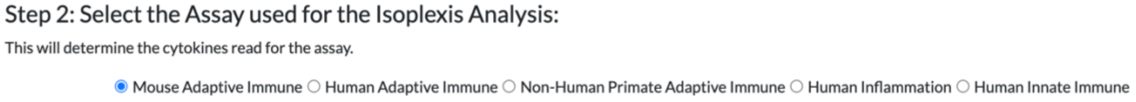


   c. Step 3: Select the conditions that will be analyzed. A drop-down menu displays the conditions that can be selected and analyzed. The order in which the conditions are selected will be the same order displayed for the visualizations in the data analysis tabs. Once the order is selected, press the "Reorder Data" button below. Note: if the user decides to select different conditions to analyze, this step, along with steps 4 and 5 must be repeated.

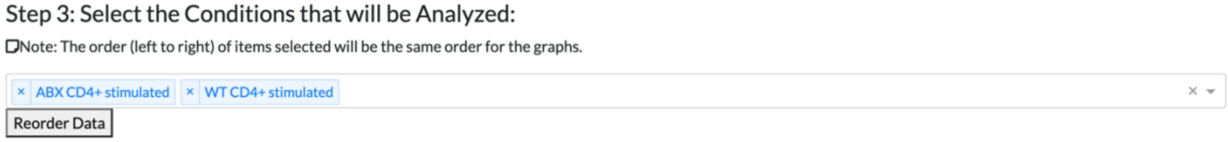


   d. Step 4: Select the "Analyze Isoplexis Data" button. Please ensure the number of cells and cytokines are correct before proceeding.

Step 4: Select the Button Below to Analyze Isoplexis Data:

Analyze Isoplexis Data

Note: If you decide to reorder your data after selecting the analysis button, you will need to repeat steps 3 and 4 again.

Please ensure that the number of cytokines and the number of cells displayed below are correct before proceeding.

If these numbers are off, double-check your original csv or excel file and also ensure the selected secretome assay is correct.

Number of Cytokines: 28

Number of Cells: 1301

e. Step 5: Individual Cytokine Analysis. Select one cytokine for the individual cytokine analysis. Note: if the user wishes to select another cytokine, step 5 must be repeated.

Step 5: To view individual cytokine analysis, select a cytokine from this list below.

Note: each analysis page has a section to view individual cytokines. If the cytokine has no value, it will not appear on this list.

Granzyme B

Analyze Individual Cytokine

In order to change individual cytokine, select desired cytokine from the dropdown menu and click 'Analyze Individual Cytokine' button again.

3. Hierarchical Clustering visualized using a dendrogram and heatmap
   a. For the clustering analysis, all or individual conditions can be selected. The visualizations will automatically update after selection.

Select subset to view for Hierarchial Clustering

This section will change both all cytokine and individual cytokine dendrograms and heatmaps.

○ All ◉ ABX CD4+ stimulated ○ WT CD4+ stimulated

   b. Dendrogram and heatmap for all cytokines. Advanced features of the visualizations include zoom, pan, auto scale, and reset axes. Visualizations can be downloaded as a png image. Hover text is also implemented in the heatmap.

Hierarchial Clustering Across All Cytokines

Dendrogram and Heatmap are clustered based on cytokine expression value similarities.

Note: the Cell ID in the legend represents the original cell ID of the uploaded data.

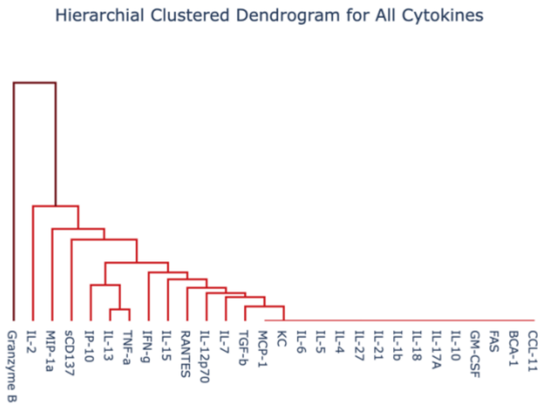


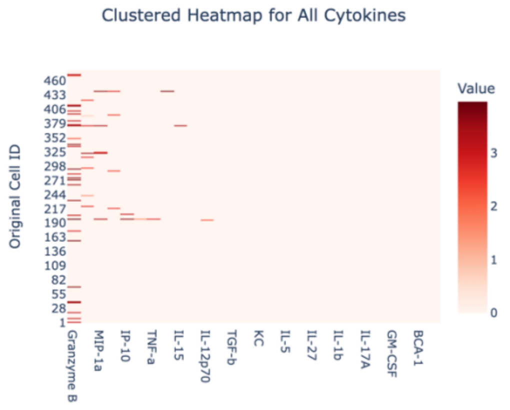

<u>Example Features (described above):</u>

Zoom

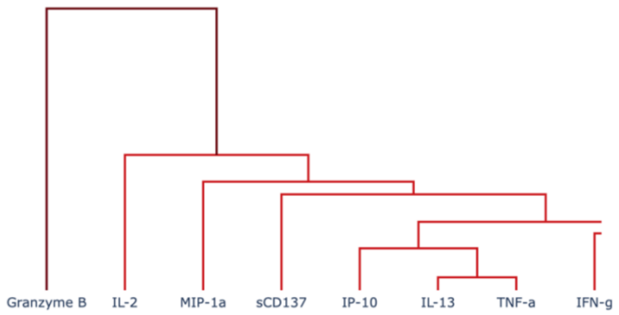


Hover text

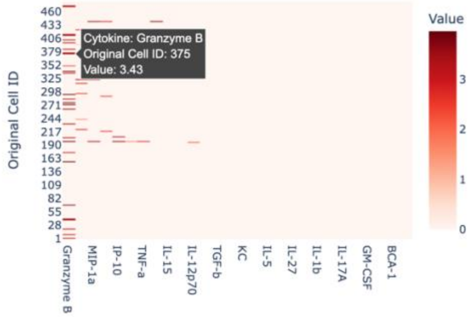


c. Dendrogram for cells with cytokine present. This analysis is based on the individual cytokine selected (2e). Only cells that contain the cytokine are used for clustering. The same features outlined above (3b) are also applicable for these visualizations.

**Hierarchial Clustering For Selected Individual Cytokine**

Dendrogram and Heatmap are clustered based on cytokine expression value similarities.

☐Note: the Cell ID in the legend represents the original cell ID of the uploaded data.

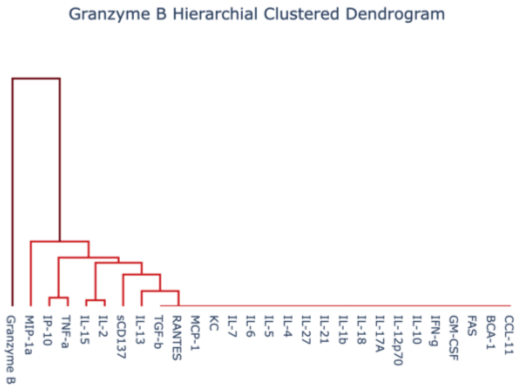


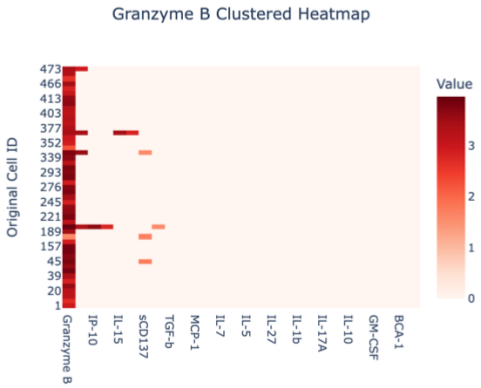


☐Note: if you would like to view individual cytokine expression of a different cytokine, repeat step 5.

4. PCA and TSNE Dimensionality Reduction

**Introduction**

PCA is a linear dimensionality reduction technique and TSNE is a non-linear dimensionality reduction technique.

☐Note: Dimensionality reduction analyses may differ from IsoSpeak. IsoSpeak uses unthresholded data, while the data file the user uploaded has a 2% threshold. Values below this threshold become 0.

TSNE might take a minute to reload since this is calculated in real-time.

a. User options for PCA and TSNE include scaling and 2D/3D visualization

**Select whether or not to normalize dimensionality reduction plots.**

Data is normally normalized before performing dimensionality reduction.

◉ Standard Scalar Normalized ○ Not Normalized

**Select to visualize the plot in 2D or 3D.**

○ 2D ◉ 3D

PCA Examples:

3D PCA

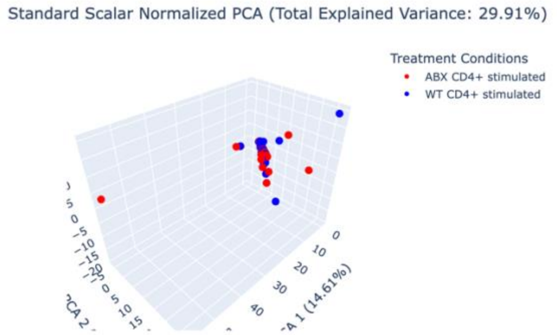


2D PCA

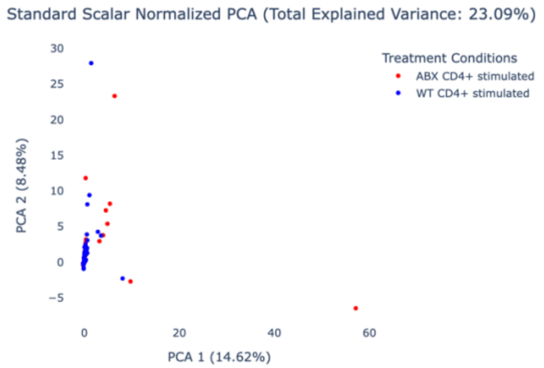


b. For TSNE, the user has the option to optimize two hyperparameters – perplexity of nearest neighbors and number of iterations.

TSNE Examples:

3D TSNE

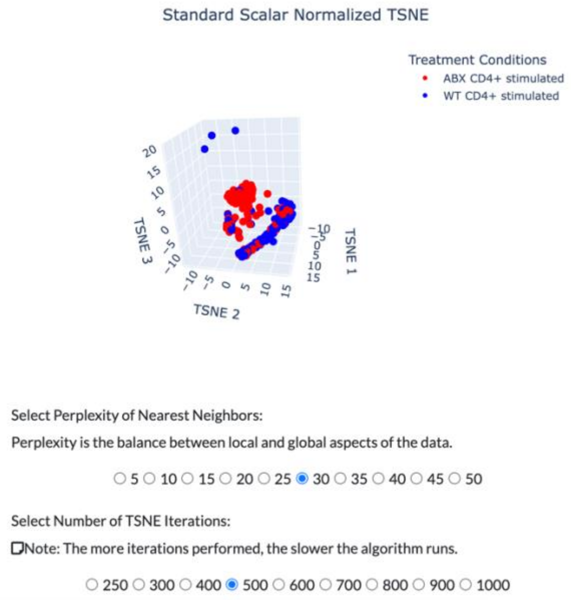


2D TSNE

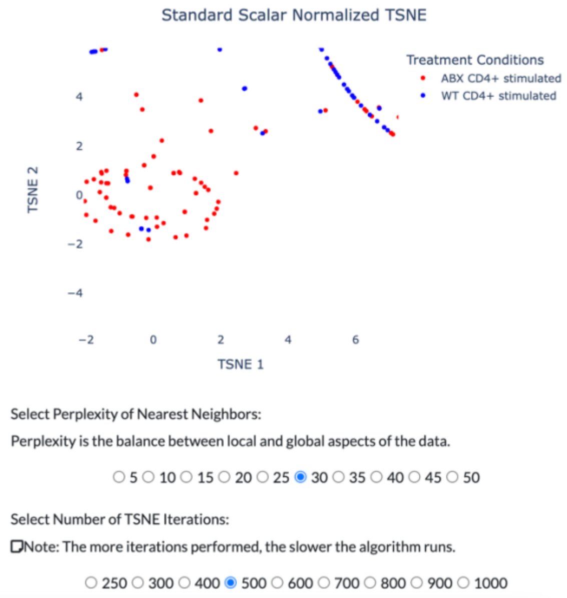


c. Advanced features of the visualizations include zoom, pan, auto scale, hover text, and reset axes. Visualizations can be downloaded as a png image. The user can also select individual treatment conditions to view.

<u>Example features (described above):</u>

Specific condition selected to view

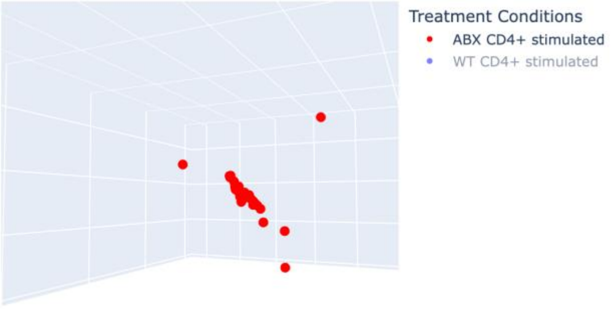


Hover text

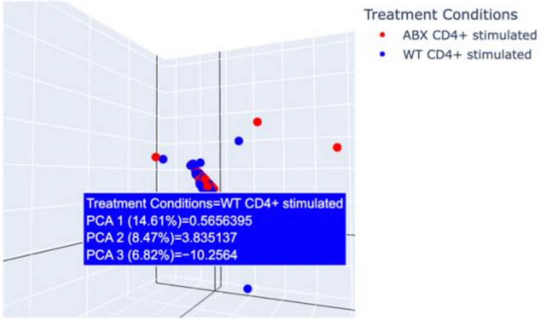


5. Polyfunctionality
   a. The number of polyfunctional cells for the assay are displayed at the top left corner of the "Polyfunctionality" tab.

### Total Number of Polyfunctional Cells: 21

   b. The stacked bar graph on the left displays the percent of cytokines secreting for each condition, which calculates the proportion of cells that express two or more proteins. The user can download these values as a CSV file.

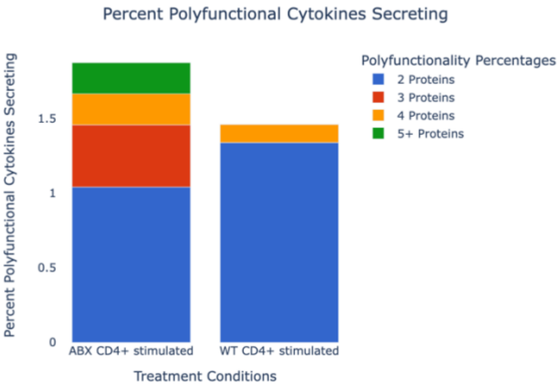


Percent Polyfunctional Cytokines Secreting: calculates the proportion of cells that express two or more proteins.

Download CSV

CSV Example Output

| Treatment Conditions | Polyfunctionality Percentages | Percent Polyfunctional Cytokines Secreting |
|---|---|---|
| ABX CD4+ stimulated | 2 Proteins | 1.041666667 |
| ABX CD4+ stimulated | 3 Proteins | 0.416666667 |
| ABX CD4+ stimulated | 4 Proteins | 0.208333333 |
| ABX CD4+ stimulated | 5+ Proteins | 0.208333333 |
| WT CD4+ stimulated | 2 Proteins | 1.339829476 |
| WT CD4+ stimulated | 3 Proteins | 0 |
| WT CD4+ stimulated | 4 Proteins | 0.12180268 |
| WT CD4+ stimulated | 5+ Proteins | 0 |

   c. The stacked bar graph on the right displays absolute abundance or proportion of dominant functional groups for the secreting cytokines as classified by Isoplexis. These values can be exported as a CSV file.

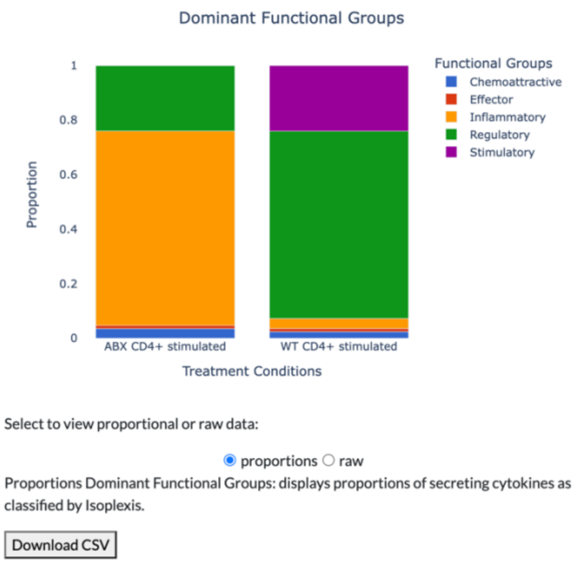


d. Advanced features of the visualizations include zoom, pan, auto scale, hover text, and reset axes. Visualizations can be downloaded as a png image. The user can also select individual treatment conditions to view.

<u>Example features (described above):</u>

Specific condition selected to view

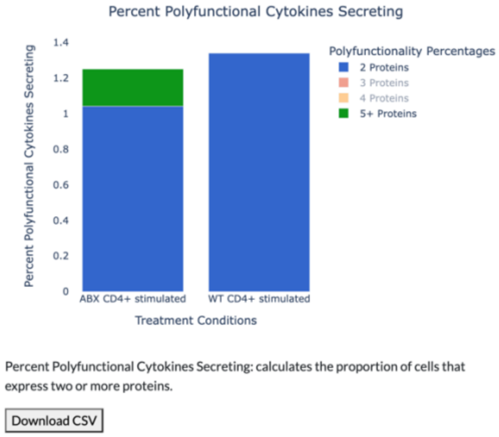


Hover text

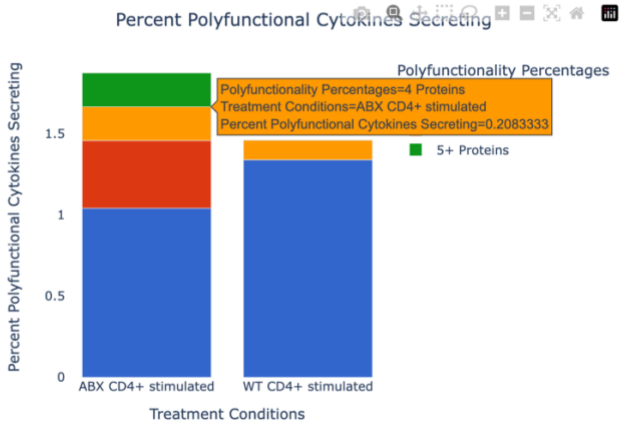


6. Distributions and Statistics
   a. Percent cytokines secreting (non-zero proportions) across all cytokines.

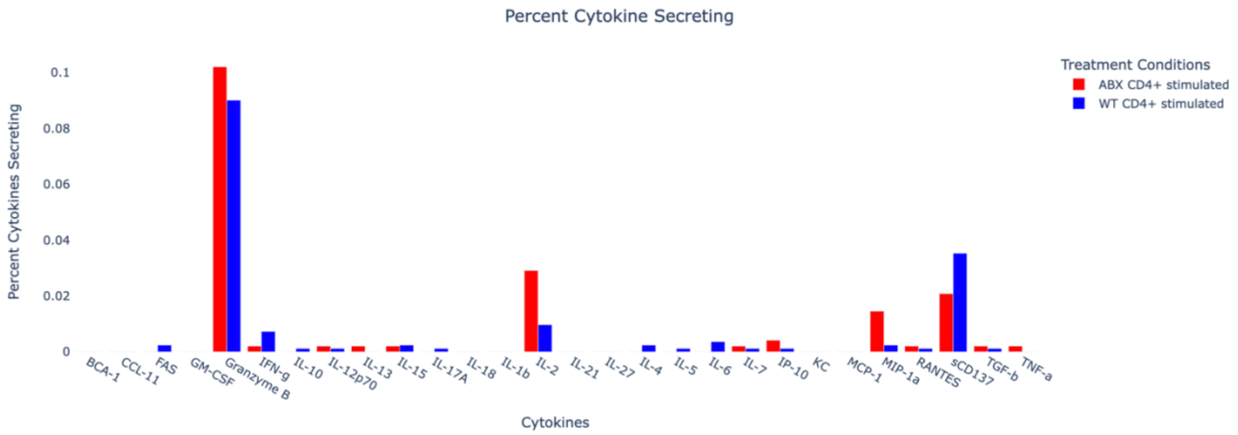


b. Individual cytokine statistical summary displays statistics for the selected cytokine (2e). The user can select all treatment conditions or an individual treatment condition.

Select Treatment Condition

◉ All ○ ABX CD4+ stimulated ○ WT CD4+ stimulated

Summary Cytokine Statistics

| Label | Value |
|---|---|
| Number of Cells with Values | 123 |
| Number of Cells with No Values | 1178 |
| Mean Across All Cells | 0.316 |
| Standard Deviation Across All Cells | 0.991 |
| Minimum Values Across All Cells | 0 |
| Maximum Values Across All Cells | 4.02 |
| Mean Across All Non-Zero Cells | 3.343 |
| Standard Deviation Across All Non-Zero Cells | 0.5131 |
| Minimum Values Across All Non-Zero Cells | 1.3 |
| Maximum Values Across All Non-Zero Cells | 4.02 |

c. Histogram of the selected cytokine (2e) and user choice of box plot, violin plot or rug plot displayed above. Additionally, the user can change the bin sizes for the histogram.

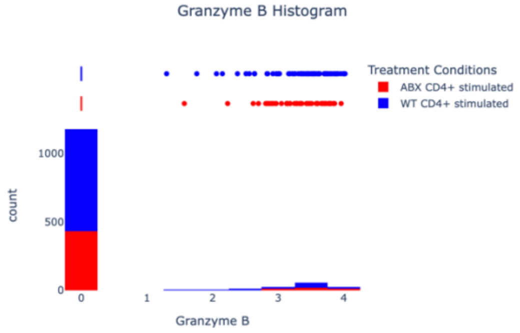


Select Bin Size:

○ 10 ◉ 20 ○ 30 ○ 40 ○ 50 ○ 60 ○ 70 ○ 80 ○ 90 ○ 100

Select Box Plot, Violin Plot or Rug Plot:

◉ Box Plot ○ Violin Plot ○ Rug Plot

d. Statistical tests include non-zero proportion and Kolmogorov-Smirnov. The user can select which conditions to compare for these tests, and the z-statistic and p-value are displayed below.

##### Statistical Tests for Differences in Cytokine Secretion

☐Note: if you would like to view individual cytokine expression of a different cytokine, repeat step 5.

*Select Condition 1:*

◉ ABX CD4+ stimulated ○ WT CD4+ stimulated

*Select Condition 2:*

○ ABX CD4+ stimulated ◉ WT CD4+ stimulated

###### Percent Cytokines Secreting - Proportion Test

The Non-Zero Proportion Test determines whether the non-zero proportion of two samples are significantly different from each other.

Non-Zero Proportion Z Statistic: 0.7108
Non-Zero Proportion P-Value: 0.4772

###### Kolmogorov-Smirnov Test

The Kolmogorov-Smirnov Test is a non-parametric test that determines if two samples are significantly different from each other.

KS-Test Proportion Z Statistic: 0.01788
KS-Test Proportion P-Value: 0.9999

e. Individual cytokine non-zero proportion bar graph is displayed on the bottom left of the “Distribution and Statistics” tab.

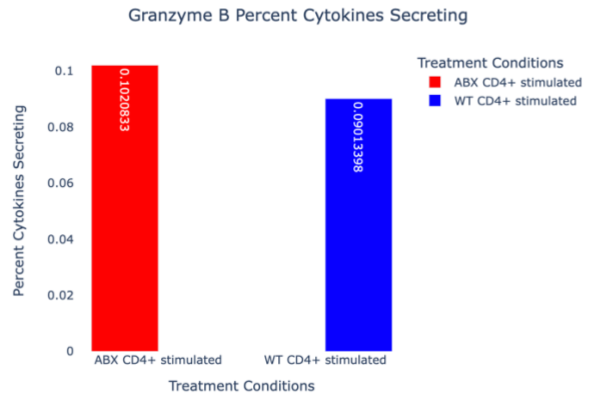


f. Individual cytokine density plot is displayed on the bottom right of the “Distribution and Statistics” tab.

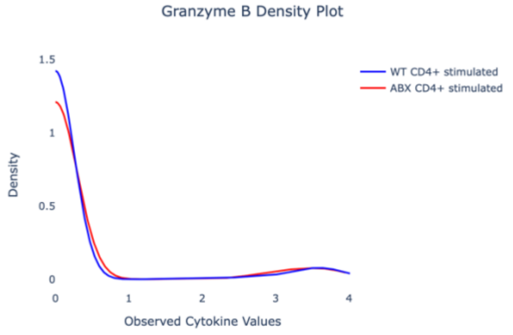


Density plots allow for the visualization of the distribution of a numeric variables for one or more groups.

g. Advanced features of the visualizations include zoom, pan, auto scale, hover text, and reset axes. Visualizations can be downloaded as a png image. The user can also select individual treatment conditions to view.

#### Section 2: FAQ

1. How often is IsoAnalytics updated?
    a. We will check for new Isoplexis single-cell assay modifications and developments monthly to ensure that our web server is keeping up with Isoplexis technology.
2. Suggestions on other methods and visualizations that you (an IsoAnalytics user) would like to see?
    a. Please reach out to Xiaowei Zhan using the. We are very open to feedback, and we would likely incorporate these suggested features/analysis/methods into our web server.
3. Data is not loading? What should you do?
    a. Check the file. Importantly, this website uses the columns "Donor", "Cell Subset" and "Stimulation" to categorize the treatment conditions. Ensure that the file has these metadata columns.

| Donor | Cell Subset | Stimulation | BCA-1 | CCL-11 |
|---|---|---|---|---|
| ABX | CD8+ | stimulated | 0 | 0 |
| ABX | CD8+ | stimulated | 0 | 0 |
| ABX | CD8+ | stimulated | 0 | 0 |
| ABX | CD8+ | stimulated | 0 | 0 |
| ABX | CD8+ | stimulated | 0 | 0 |
| ABX | CD8+ | stimulated | 0 | 0 |
| ABX | CD8+ | stimulated | 0 | 0 |
| ABX | CD8+ | stimulated | 0 | 0 |
| ABX | CD8+ | stimulated | 0 | 0 |
| ABX | CD8+ | stimulated | 0 | 0 |
| ABX | CD8+ | stimulated | 0 | 0 |
| ABX | CD8+ | stimulated | 0 | 0 |

    b. Check that the correct assay on the "Upload" tab has been selected. Currently there are five single-cell assays (listed below). This website relies on the presence of the correct list of cytokines used for each assay.

**Step 2: Select the Assay used for the Isoplexis Analysis:**

This will determine the cytokines read for the assay.

◉ Mouse Adaptive Immune ○ Human Adaptive Immune ○ Non-Human Primate Adaptive Immune ○ Human Inflammation ○ Human Innate Immune

    c. Still not working? Please contact Xiaowei Zhan using the. We will help you troubleshoot.
4. Notice a bug or glitch?
    a. For issues, please create a new issue through https://github.com/suziepalmer10/Isoplexis_Data_Analysis/issues.
